# *Babesia duncani* as a model organism to study the development, virulence and drug susceptibility of intraerythrocytic parasites *in vitro* and *in vivo*

**DOI:** 10.1101/2021.12.01.470698

**Authors:** Anasuya C. Pal, Isaline Renard, Pallavi Singh, Pratap Vydyam, Joy E. Chiu, Sovitj Pou, Rolf W. Winter, Rozalia Dodean, Lisa Frueh, Aaron C. Nilsen, Michael K. Riscoe, J. Stone Doggett, Choukri Ben Mamoun

## Abstract

Hematozoa are a subclass of protozoan parasites that invade and develop within vertebrate red blood cells to cause the pathological symptoms associated with diseases of both medical and veterinary importance such as malaria and babesiosis. A major limitation in the study of the most prominent hematozoa, *Plasmodium spp,* the causative agents of malaria, is the lack of a broadly accessible mouse model to evaluate parasite infection *in vivo* as is the case for *P. falciparum* or altogether the lack of an *in vitro* culture and mouse models as is the case for *P. vivax*, *P. malariae* and *P. ovale*. Similarly, no *in vitro* culture system exists for *Babesia microti*, the predominant agent of human babesiosis. In this study, we show that human red blood cells infected with the human pathogen *Babesia duncani* continuously propagated in culture, as well as merozoites purified from parasite cultures, can cause lethal infection in immunocompetent C3H/HeJ mice. Furthermore, highly reproducible parasitemia and survival outcomes were established using specific parasite loads and different mouse genetic backgrounds. Using the combined in culture-in mouse (ICIM) model of *B. duncani* infection, we demonstrate that current recommended combination therapies for the treatment of human babesiosis, while synergistic in cell culture, have weak potency *in vitro* and failed to clear infection or prevent death in mice. Interestingly, using the ICIM model, we identified two new endochin-like quinolone prodrugs, ELQ-331 and ELQ-468, that alone or in combination with atovaquone are highly efficacious against *B. duncani* and *B. microti*. The novelty, ease of use and scalability of the *B. duncani* ICIM dual model make it an ideal system to study intraerythrocytic parasitism by protozoa, unravel the molecular mechanisms underlying parasite virulence and pathogenesis, and accelerate the development of innovative therapeutic strategies that could be translated to unculturable parasites and important pathogens for which an animal model is lacking.

**Author Summary:** Use of model organisms is vital to the understanding of virulence and pathogenesis of a large number of human and animal pathogens. In case of hematozoan parasites that invade and develop within vertebrate erythrocytes, the studies are challenging because of the dearth of small animal model systems and the lack of continuous parasite growth in *in vitro* culture conditions. Here, we report a small animal model of lethal infection of *Babesia duncani*, one of the causative agents of human babesiosis. We show that *in vitro* cultured parasites and as well as parasites propagated *in vivo* can establish highly reproducible parasitemia which is dependent on the parasite load and also defined by different mouse genetic backgrounds. We further use this combined in culture-in mouse (ICIM) model of *B. duncani* infection to demonstrate the anti-babesial efficacy of two novel endochin like quinoline compounds. We propose that this ICIM dual model of *B. duncani* is an ideal system to get insights into protozoan intraerythrocytic parasitism, virulence, pathogenesis, and therapeutics and will open the vista to other important pathogens which are unculturable or lack an animal model.

## Introduction

The Apicomplexa phylum encompasses a diverse group of mostly obligate intracellular parasites that cause a number of diseases in humans and animals. A subclass of these parasites includes *Plasmodium* and *Babesia* parasites, the causative agents of malaria and babesiosis, respectively, which are hematozoan pathogens that invade and replicate within host red blood cells to produce a varying number of daughter parasites also known as merozoites [1]. Each merozoite develops within the host erythrocyte by consuming a large set of host nutrients and undergoes various cycles of asexual replication to produce new merozoites following destruction of the host cell. These cycles of invasion, replication and rupture can lead to major pathology and often death as in the case of human and avian malaria, as well as human and bovine babesiosis [2, 3]. Because of the major health and economic impact of these diseases, efforts are required to control the development and transmission of the pathogens involved, in order to eliminate these diseases. Such efforts rely on the development of effective control measures and therapeutic strategies, which themselves require a thorough understanding of key biological clues about pathogen metabolic needs, mode of development and transmission, determinants of virulence, and susceptibility to novel drugs. In the case of the human malaria parasite *Plasmodium falciparum*, this knowledge reached an inflection point following the establishment of a continuous and long-term *in vitro* culture system in 1976 by Trager and Jensen [4]. Unfortunately, to this date, no small animal model is available to recapitulate the development, replication and differentiation of *P. falciparum* in mice. Alternative models include humanized mice, which are highly immunocompromised and transfused with human red blood cells or hematopoietic progenitor cells and subsequently infected with adapted *P. falciparum* strains [5]. Such models have been used to evaluate short-term efficacy of drugs targeting asexual replication and/or sexual differentiation [6]. However, these models are not cost-effective, only support short blood stage developmental cycles and are not amenable to studies aimed to elucidate mechanisms of virulence, pathogenesis and host-pathogen interactions. For other human malaria parasites such as *P. vivax*, *P. ovale* and *P. malaria,* there are neither *in vitro* culture systems nor mouse models of infection, thereby hindering the study of these most prominent hematozoan pathogens. Therefore, most of the existing knowledge about *in vivo* intraerythrocytic *Plasmodium* development, immunity and protection relies on knowledge gained from studies of murine malaria parasites *P. yoelii* and *P. berghei* [7, 8]. The biology of these models, however, does not necessarily translate to that of human parasites [7, 9–11].

*Babesia* parasites are phylogenetically closely related to *Plasmodium* species and cause a malaria-like illness, which in susceptible individuals can lead to hemolytic anemia, acute respiratory distress syndrome, hepatosplenomegaly, multiorgan failure and possibly death [12]. Human babesiosis is an emerging tick-borne disease and is spreading rapidly mostly due to changes in the geographic distribution of the vector, anthropogenic factors and global warming [1, 12, 13]. Several species of *Babesia,* including *B. microti*, *B. duncani* and *B. divergens*, are known to cause infection in humans [1]. *B. microti*, the most commonly reported *Babesia* pathogen, can be propagated in immunocompetent and immunocompromised mice but has not yet been successfully maintained continuously *in vitro* in human red blood cells [1]. Conversely, *Babesia divergens* can be successfully propagated *in vitro* in human red blood cells but so far no mouse model exists for this parasite. The *B. duncani* WA-1 isolate [14] was successfully cultured in hamster red blood cells [15] and more recently continuously propagated in both hamster and human red blood cells [16]. The continuous culture of the parasite in human red blood cells made it possible to screen chemical libraries to evaluate the efficacy of new drugs, study drug-drug interactions and probe the mode of action of active compounds [17, 18]. Interestingly, the WA-1 isolate was also successfully propagated in mice and hamsters following intraperitoneal administration. In these models, parasite inoculation was followed by a rapid increase in parasitemia and severe pathology [15, 19, 20].

Here we report an optimized *B. duncani* ICIM (in culture-in mouse) model that combines *in vitro* cell culture of the parasite in human red blood cells with a highly precise model of lethal infection in mice. Using this model, we show that current clinically approved drug combinations while synergistic in culture have limited potency *in vitro* and in mice. Furthermore, we report the successful use of this model in the discovery of two new endochin-like quinolones with potent antibabesial activity *in vitro* and strong efficacy in mice against *B. duncani* as well as *B. microti*.

## Results

### Establishment of an in culture-in mouse (ICIM) model of *B. duncani* **infection**

Continuous *in vitro* culture of *B. duncani* WA-1 isolate in human red blood cells was previously reported using commercially available HL-1 and Claycomb media [16]. In this system, the different development stages of the parasite (young ring, mature ring, filamentous, young tetrad and mature tetrad) formed during asexual replication are typically observed (Fig 1A), with a replication time of 22-24 h and parasitemia levels reaching as high as 20% [16]. As shown in Fig 1B, parasitemia levels can also be evaluated using SYBR Green I-based assays. To determine whether *in vitro*-cultured parasites can initiate infection in mice, female C3H/HeJ mice (n = 3 mice/group) were injected intravenously (IV) with either 8.5 x 10^5^ *B. duncani*-infected hRBCs or 2 x 10^7^ cell-free merozoites purified from *in vitro B. duncani* culture (Fig 1C). As shown in Figure 1D, parasitemia was established in both groups of mice by 8 and 12 days post-infection (DPI) following the administration of infected hRBCs and cell-free merozoites, respectively. In both groups, peak parasitemia reached 4-7% and the animals became sick and had to be euthanized by DPI 11 and DPI 15, depending on whether they received infected hRBCs or cell-free merozoites, respectively (Fig 1E).

**Fig 1.**
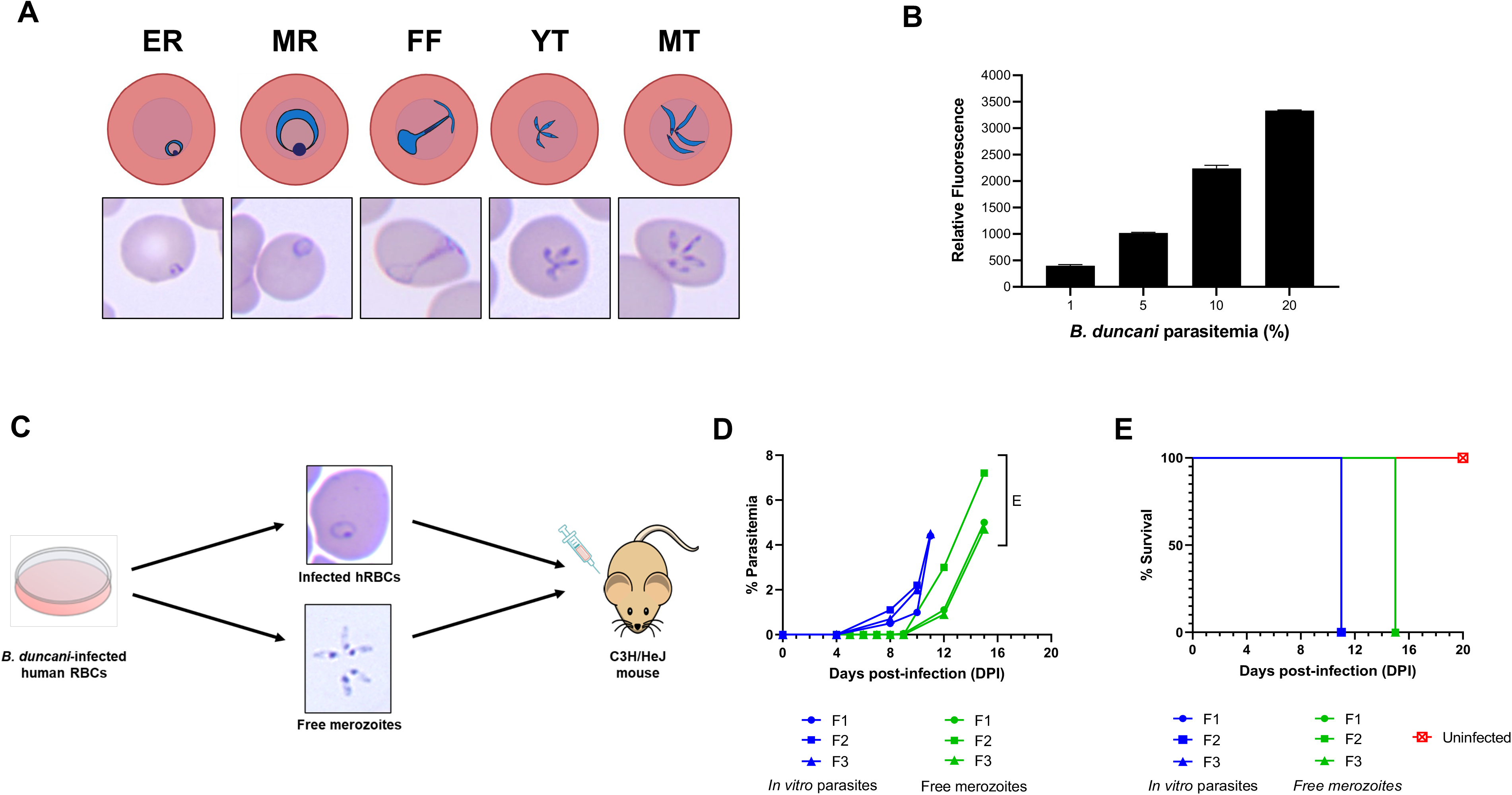
Continuous *in vitro* culture of *B. duncani* in human RBCs and transmission of cultured parasites to mice. **A)** Schematic representation of different stages of *B. duncani* in *in vitro* culture and corresponding Giemsa-stained slides of parasitized hRBCs (ER = early ring; MR = mature ring; FF = filamentous form; YT = young tetrad; MT = mature tetrad). **B)** Correspondence between fluorescence reading using SYBR Green I staining (497/520 nm) and percent parasitemia determined by light microscopy examination of Giemsa-stained thin blood smears. **C)** Schematic representation of transmission of *in vitro*-cultured *B. duncani* parasites or purified merozoites into C3H/HeJ mice. **D)** Parasitemia profile over time in female C3H/HeJ mice (n = 3 mice/group) following infection with *in vitro* parasites (blue) or purified merozoites (green). E: euthanized. **E)** Kaplan-Meier plot of percent survival of uninfected mice (red), mice infected with *in vitro* parasites (blue) or mice infected with purified merozoites (green).

While the transmission of *B. duncani* infection from *in vitro* to *in vivo* provides a unique opportunity to manipulate the parasite and evaluate the impact of such manipulations on virulence and pathogenesis, further optimization of *B. duncani* infection in mice was required in order to establish a reliable *in vivo* model. C3H/HeJ were inoculated IV with parasite loads ranging from 10^7^ to 10^2^ *B. duncani*-infected red blood cells (iRBCs) collected from a previously infected mouse. As shown in Fig 2A, consistent infection in both male (n = 3 mice/infection dose) and female (n = 3 mice/infection dose) mice was established in a dose-dependent manner. All infections resulted in lethal outcome with a severity proportional to the administered dose. Mice inoculated with 10^7^ and 10^6^ *B. duncani*-iRBCs showed an acute increase in parasitemia (up to 35%) within a short span of time (DPI 3-5), became moribund and had to be euthanized by DPI 5-7 (Fig 2). Animals infected with doses between 10^5^ and 10^2^ iRBCs showed a delayed onset of infection with a lower peak parasitemia compared to those administered with 10^7^/10^6^ iRBCs, proportional to the administered dose (Fig 2A). The parasite burden was also function of the infection dose, where animals infected with 10^5^ iRBCs showed a peak parasitemia up to 9%, whereas mice infected with 10^2^ iRBCs reached no more than 2.5% (Fig 2A). Nonetheless, all mice in every infection group became equally sick and had to be euthanized (Fig 2B). Interestingly, parasitemia levels appeared consistently higher in female mice than in male mice (Fig 2A), despite the same endpoint (Fig 2B).

**Fig 2.**
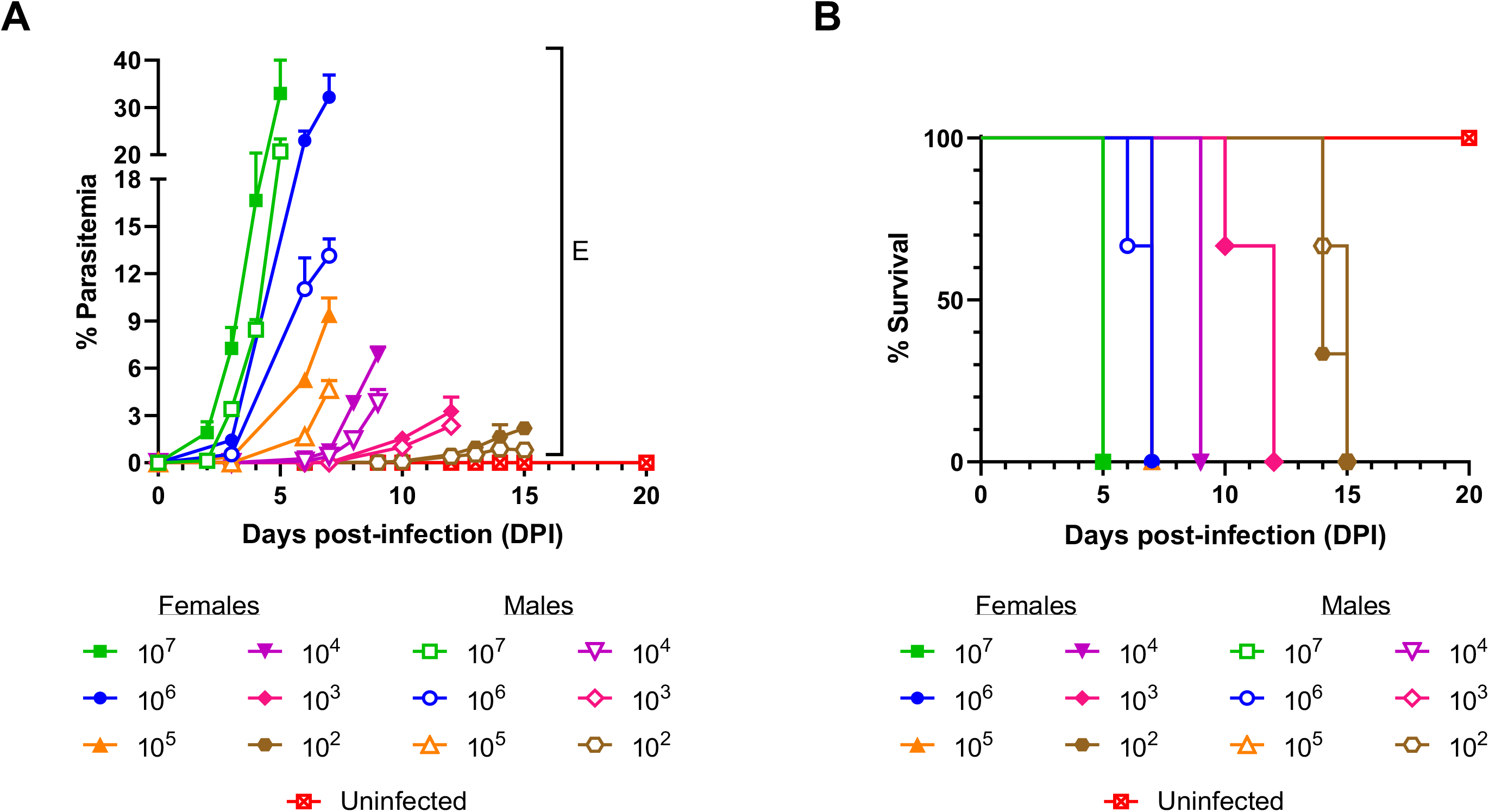
Lethal *B. duncani* infection model in immunocompetent mice. **A)** Parasitemia profile of *B. duncani* infection in female (solid symbols) and male (open symbols) C3H/HeJ mice infected IV with doses ranging from 107 to 102 iRBCs. E: Euthanized. **B)** Kaplan-Meier plot of percent survival of female (solid symbols) and male (open symbols) C3H/HeJ mice infected with different doses of *B. duncani*. Uninfected mice (red) were used as control.

### Influence of mouse genetic background and route of administration on the response to *B. duncani* infection

We explored the importance of the host genetic background, as well as the route used for parasite administration, on the susceptibility of mice to *B. duncani* infection. C3H/HeJ, Balb/cJ and C57BL/6J mice (n = 6 mice/group; 3 females and 3 males) were infected IV or intraperitoneally (IP) with 10^4^ *B. duncani*-iRBCs. As shown in Fig 3, all C3H/HeJ mice, inoculated by either IV or IP route, showed detectable parasitemia by DPI 6 or 7, with a peak parasitemia of 6% for males and 10.7% for females, respectively (Fig 3A and 3D). All mice became sick and had to be euthanized by DPI 10 (Fig 3G and 3H). In contrast, C57BL/6J mice appeared resistant to *B. duncani* infection and showed no detectable parasitemia in thin blood smears throughout the 24-day monitoring period (Fig 3C and 3F), resulting in 100% survival (Fig 3G and 3H). Previous studies have reported successful *B. duncani* infection in C57BL/6J. However, the dose administered was much higher than the dose used in the present study [21]. Interestingly, Balb/cJ mice showed intermediate range of susceptibility with 40-60% survival. Some of the IV-infected Balb/cJ mice developed parasitemia between 3-9% by DPI 9, became sick and had to be euthanized; whereas the rest of the cohort cleared the infection (Fig 3B). The IP-infected Balb/cJ mice showed a similar profile but reached lower levels of peak parasitemia (1-2%) (Fig 3E).

**Fig 3.**
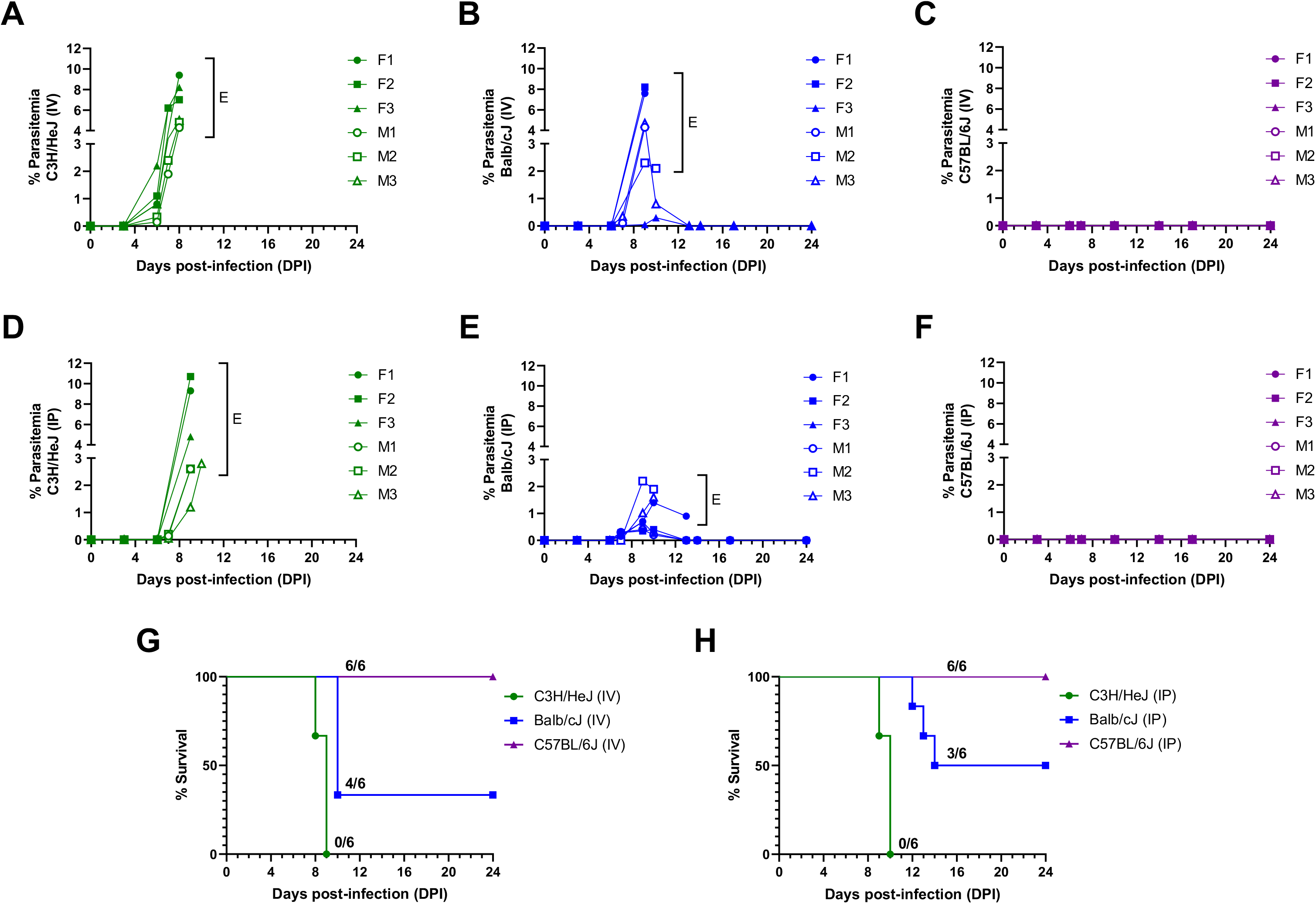
Genetic background of mice defines *B. duncani* infection outcome. Immunocompetent female and male mice from three genetic backgrounds were infected with 104 *B. duncani*-infected RBCs by either intravenous (IV) or intraperitoneal (IP) route and parasitemia was monitored over time. **A-C)** Evolution of parasitemia over time following IV administration of iRBCs in (**A**) C3H/HeJ, (**B**) Balb/cJ and (**C**) C57BL/6J mice. **D-F)** Evolution of parasitemia over time following IP administration of iRBCs in (**D**) C3H/HeJ, (**E**) Balb/cJ and (**F**) C57BL/6J mice. Female and male profiles are shown in solid and open symbols, respectively. E: euthanized. **G-H)** Kaplan-Meier plot showing percent survival of C3H/HeJ, Balb/cJ, and C57BL/6J mice post (**G**) IV or (**H**) IP infection with *B. duncani*-iRBC.

### Role of host immunity on the susceptibility to *B. duncani* infection

To assess whether the resistance of C57BL/6J mice was the result of rapid clearance by the immune system or inability of the parasite to infect RBCs from this genetic background at the selected 10^4^ *B. duncani*-iRBCs, we compared the parasitemia profile between immunocompetent and immunocompromised mice following infection with *B. duncani*. SCID (severe combined immunodeficient) mice from C3H/HeJ, Balb/cJ and C57BL/6J backgrounds (C3H/HeJ-SCID, Balb/cJ-SCID and C57BL/6J-SCID) were infected IV with 10^4^ *B. duncani*-iRBCs and their parasite load and health status were monitored over time. The C3H/HeJ-SCID mice reached a higher parasitemia (up to 60%) than their wild-type (WT) counterpart (maximum load of 10%) (Fig 4A and 3A) and had to be euthanized by DPI 14 (Fig 4F). The difference in peak parasitemia between WT and SCID mice was also observed in the case of Balb/cJ mice (8% *vs*. 60% for Balb/cJ and Balb/cJ-SCID, respectively) (Fig 3B and 4B) and all but one Balb/cJ-SCID mouse had to be euthanized by DPI 17 (Fig 4F). One Balb/cJ-SCID mouse did not reach high parasitemia (peak around 6%) and cleared the infection, suggesting a leakiness in the *scid* mutation. Interestingly, in contrast to the C57BL/6J immunocompetent mice (Fig 3C), all C57BL/6J-SCID mice developed high parasitemia (up to 40%) and reached the endpoint by DPI 22 (Fig 4C and 4F). Together these data indicate that susceptibility to *B. duncani* infection is strongly linked to the immune status of the host.

**Fig 4.**
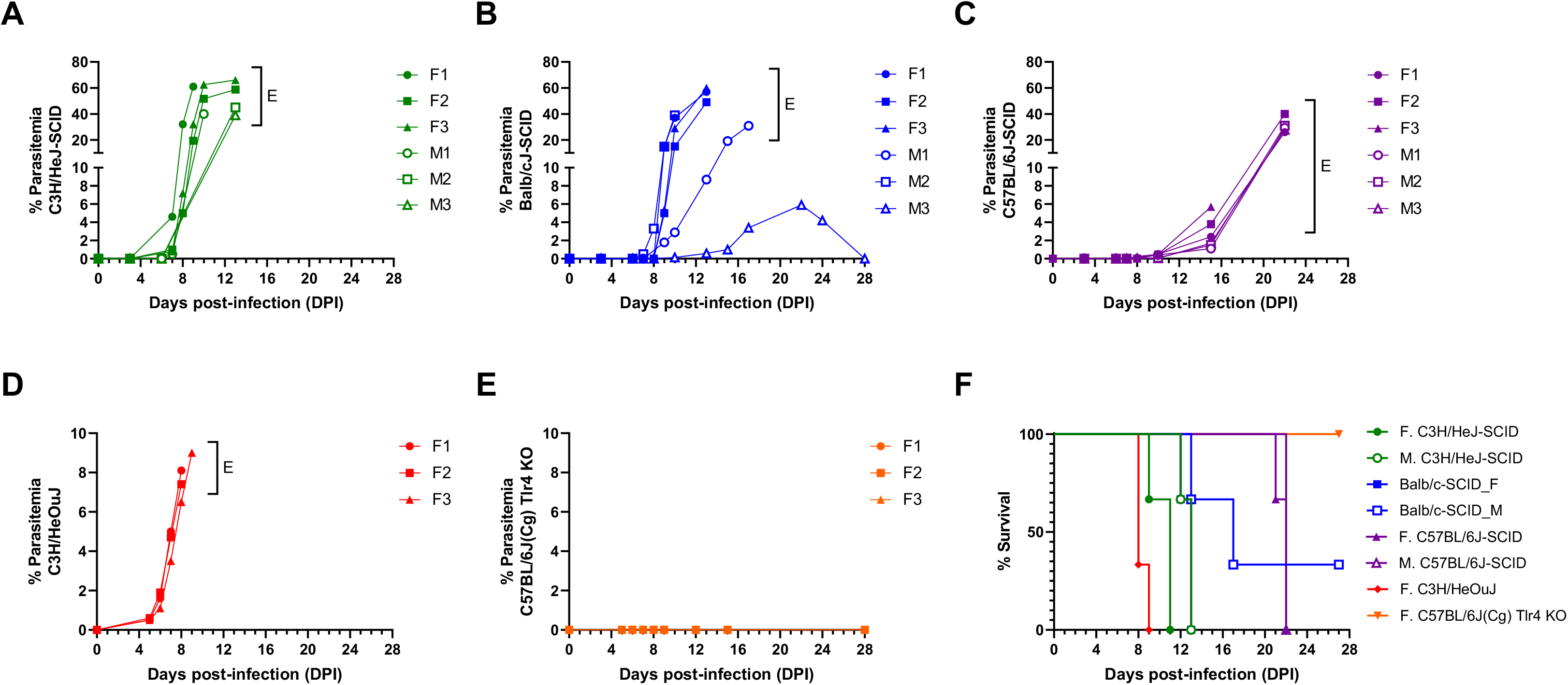
Role of host immunity against *B. duncani* infection *in vivo*. Female C3H/HeJ-SCID, Balb/cJ-SCID, C57BL/6J-SCID, C3H/HeOuJ, or C57BL/6J(Cg) *Tlr4* KO mice were infected IV with 104 *B. duncani*-iRBCs. **A-E)** Evolution of parasitemia over time in (**A**) C3H/HeJ-SCID, (**B**) Balb/cJ-SCID, (**C**) C57BL/6J-SCID, (**D**) C3H/HeOuJ and (**E**) C57BL/6J(Cg) *Tlr4* KO. Female and male profiles are shown in solid and open symbols, respectively. E: euthanized. **F)** Kaplan- Meier plot showing percent survival of the above-mentioned groups of mice post IV infection with *B. duncani*-iRBC.

Since the highly susceptible immunocompetent C3H/HeJ mouse is known to carry a missense mutation in the 3^rd^ exon of the Toll-like receptor TLR4 gene [22], we investigated whether *Tlr4* plays a role in the susceptibility to *B. duncani* infection. We infected C3H/HeOuJ mice that possess a WT *Tlr4* and a *Tlr4* KO strain on C57BL/6 background (C57BL/6J(Cg) *Tlr4* KO) with 10^4^ *B. duncani*-iRBC and monitored parasitemia and mouse survival post-infection. We found that the C3H/HeOuJ mice were equally susceptible to *B. duncani* infection as their C3H/HeJ counterpart (Fig 4D and 4F). Similarly, C57BL/6J(Cg) *Tlr4* KO mice were found equally resistant to the infection as the C57BL/6J mice (Fig 4E and 4F), indicating that *Tlr4* does not play a role in host susceptibility to *B. duncani* infection.

### Efficacy of clinically approved drug combinations against *B. duncani* **growth *in vitro and in vivo***

With an optimized *in vivo* mouse model for *B. duncani* propagation, we next attempted to correlate the *in vitro* and *in vivo* efficacy of established antibabesial drugs. Our previous work showed that, when used individually, the clinically approved antibabesial drugs (azithromycin, atovaquone, quinine and clindamycin), except for atovaquone, have weak *in vitro* activity against*B. duncani* [16]. Here, using the *in vitro* culture system of *B. duncani* in human RBCs, we performed drug-drug interaction studies to assess the possible additive or synergistic effect between the clinical combinations: azithromycin + atovaquone and quinine + clindamycin. As shown in Fig 5A and 5B, combinations of azithromycin + atovaquone and quinine + clindamycin were found to be synergistic with mean fractional inhibitory concentrations (ƩFIC50) of 0.6 (Table 1 and Fi.g 5). The efficacy of these drug combinations was further evaluated *in vivo* in a mouse model of *B. duncani* infection. Female C3H/HeJ mice (n = 5 mice/group) were infected with 10^3^ *B.duncani*-infected RBCs and treated with either azithromycin + atovaquone (10 + 10 mg/kg), quinine + clindamycin (10 + 10 mg/kg) or the vehicle alone (PEG-400) for 10 days starting at DPI 1. Interestingly, only the azithromycin + atovaquone combination showed partial efficacy at controlling the infection with two out of five animals in this treatment group not showing any parasitemia and surviving for the entire duration of study (DPI 45) (Fig 5C and 5D). The other three mice in this group started showing parasitemia from *ca.* DPI 20, became sick and were euthanized by DPI 25, with a parasite load ranging from 2-4%. All mice in the quinine + clindamycin treatment group, as well as the vehicle-treated control group, showed increasing parasitemia from DPI 8, reached up to 6% parasitemia burden between DPI 10-14 when they became moribund and had to be euthanized (Fig 5C and 5D).

**Fig 5.**
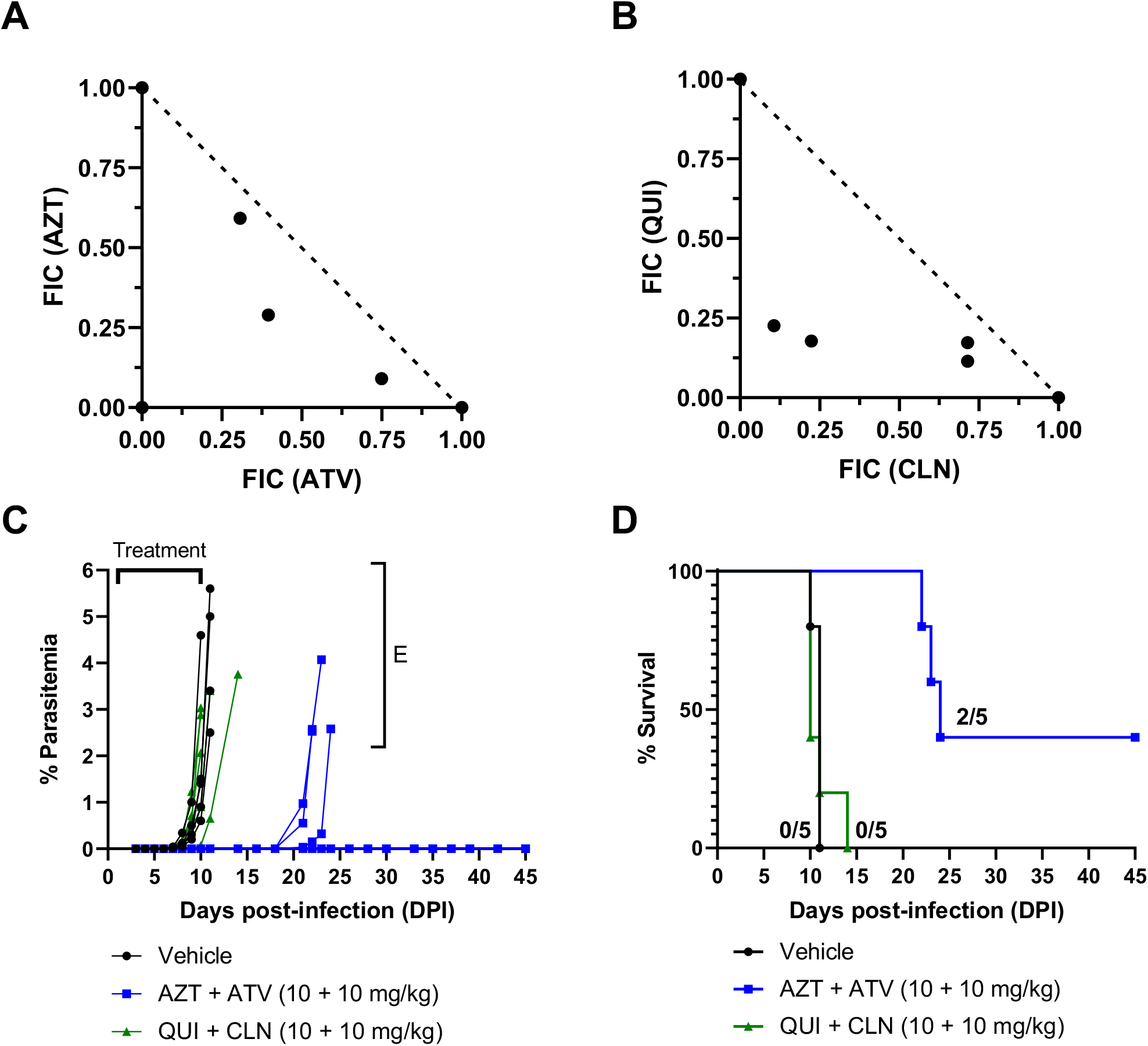
Efficacy of clinically approved drug combinations against *B. duncani* parasites *in vitro* and *in vivo*. **A-B)** Isobolograms showing the synergistic interaction between (**A**) azithromycin (AZT) and atovaquone (ATV) and (**B**) quinine (QUI) and clindamycin (CLN). The dotted line plotted between the individual mean FIC50 values of each drug serves as the additive line. FIC: fractional inhibitory concentration. **C)** Parasitemia profile of *B. duncani*-infected female C3H/HeJ mice following treatment with vehicle (PEG-400) alone, azithromycin + atovaquone (AZT + ATV) or quinine + clindamycin (QUI +CLN) at 10 + 10 mg/kg. Treatment was administered daily for 10 days (DPI 1-10). E: euthanized. **D)** Kaplan-Meier plot showing percent survival of *B. duncani*-infected mice treated with vehicle (PEG 400), azithromycin + atovaquone (AZT + ATV) or quinine + clindamycin (QUI +CLN).

**Table 1.** Evidence for synergistic activity of azithromycin and atovaquone, quinine and clindamycin, ELQ-331 and atovaquone and ELQ-468 and atovaquone against *B. duncani in vitro*.

| Drug combinations | Mean ΣFIC<sub>50</sub> | Type of Interaction |
| --- | --- | --- |
| Azithromycin + Atovaquone | 0.6 | Synergistic |
| Quinine + Clindamycin | 0.6 | Synergistic |
| ELQ-331 + Atovaquone | 0.6 | Synergistic |
| ELQ-468 + Atovaquone | 0.8 | Synergistic |

### ELQ compounds demonstrate *in vitro* and *in vivo* efficacy against *B. duncani*

Partial effectiveness of the clinically approved drug combinations against *B. duncani* infection (Fig 5C and 5D), together with the fact that these treatments are frequently associated with severe side effects, newer and safer therapeutic approaches are needed. Using our optimized *B. duncani* infection ICIM model, we therefore tested the efficacy of two endochin-like quinolones (ELQs). We have recently used the *in vitro* culture system to conduct Structure-Activity Relationship (SAR) studies on ELQ derivatives and identified ELQ-502 as a potent antiparasitic drug [17]. During this screening, two additional prodrugs, ELQ-331 and ELQ-468 (Fig 6A) were identified as potent inhibitors of *B. duncani* growth *in vitro* with IC50 values of 15 nM and 141 nM, respectively (Fig 6B and 6C) [17]. *In vitro* evaluation of ELQ-300 and ELQ-446, the parent drugs of ELQ-331 and ELQ-468, respectively, resulted in low nanomolar inhibition of *B. duncani*, with IC50 values of 36 and 34 nM, respectively (Fig 6A-C). In addition to their high efficacy against *B. duncani*, the low toxicity of ELQ-331 and ELQ-468 in mammalian cells resulted in highly favorable therapeutic indices (Table 2). Furthermore, *in vitro* combination studies of ELQ-331 and ELQ-468 with atovaquone identified synergistic interactions between these ELQ derivatives and atovaquone with mean fractional inhibitory concentrations (ƩFIC50) of 0.6 and 0.8 for ELQ-331 + atovaquone and ELQ-468 + atovaquone, respectively (Fig 6D and 6E and Table 1). These results are consistent with previous evaluation of other ELQs with atovaquone [17, 23].

**Fig 6.**
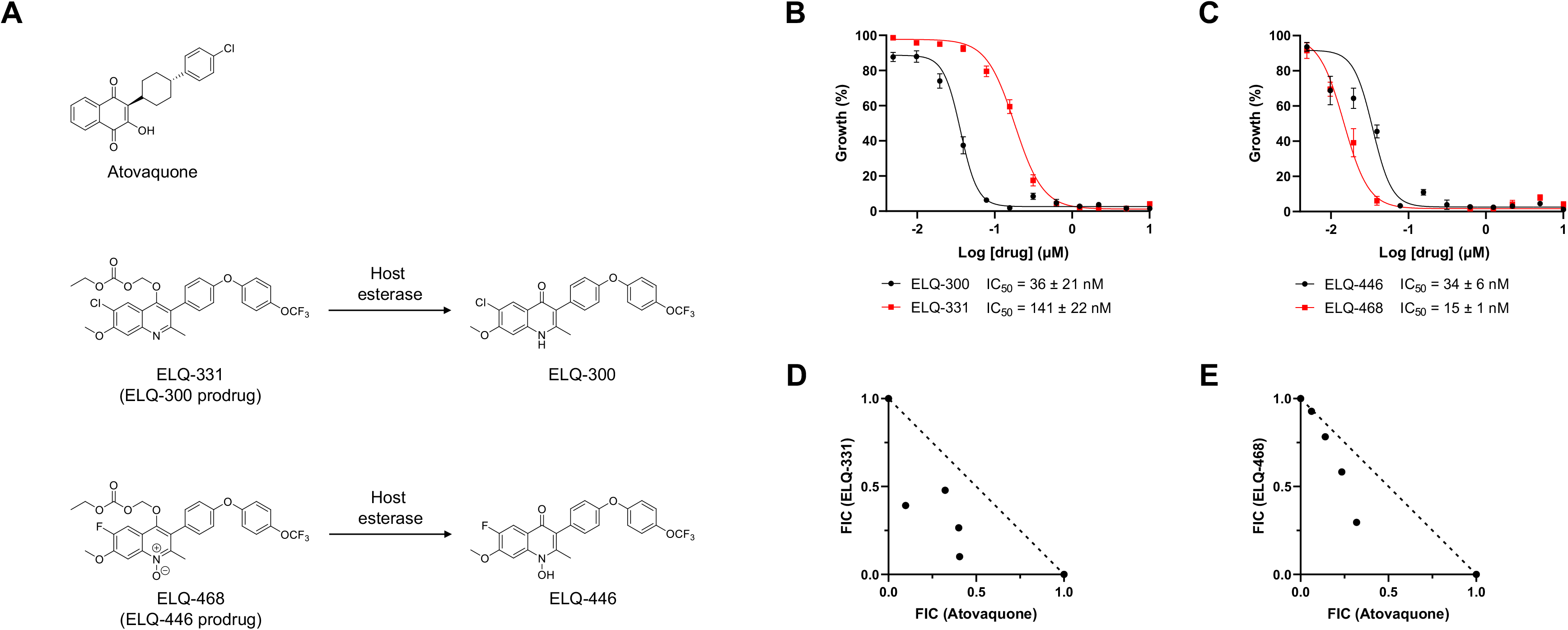
*In vitro* evaluation of ELQs against *B. duncani*. **A)** Chemical structures of Atovaquone, ELQ-331 and its parent drug, ELQ-300 and ELQ-468 and its parent drug, ELQ-446. The prodrugs are cleaved by host esterase to release the parent compound. **B-C)** Dose-response curves of (**B**) ELQ-300 and ELQ-331 and (**C**) ELQ-446 and ELQ-468 and corresponding IC50 values. **D-E)** Isobolograms showing synergistic interactions between (**D**) ELQ-331 and atovaquone and (**E**) ELQ-468 and atovaquone. The dotted line plotted between the individual mean FIC50 values of each drug serves as the additive line. FIC: fractional inhibitory concentration.

**Table 2.** Activity of ELQ-331 and ELQ-468 in *B. duncani*-infected human erythrocytes, HeLa, HepG2, HEK293 and hTERT cells.

| Prodrug | $\text{IC}_{50}$ | | | | | Therapeutic Index |
| --- | --- | --- | --- | --- | --- | --- |
|  | *B. duncani* | HeLa | HepG2 | HEK293 | hTERT |  |
| ELQ-331 | 141 ± 22 nM | > 10 $\mu\text{M}$ | > 10 $\mu\text{M}$ | > 10 $\mu\text{M}$ | > 10 $\mu\text{M}$ | > 71 |
| ELQ-468 | 15 ± 1 nM | > 10 $\mu\text{M}$ | > 10 $\mu\text{M}$ | > 10 $\mu\text{M}$ | > 10 $\mu\text{M}$ | > 667 |

ELQ-331 and ELQ-468 were further tested against *B. duncani in vivo*. Female C3H/HeJ mice (n = 5-10 mice/group) were infected with 10^3^ *B. duncani*-infected iRBC and monotherapies using ELQ-331, ELQ-468 or atovaquone or combination therapies with ELQ-331+ atovaquone (10 + 10 mg/kg) or ELQ-468 + atovaquone (10 + 10 mg/kg) were administered for 10 days (DPI 1-10). As shown in Fig 7A and 7B, animals treated with ELQ-331 and ELQ-468, either as monotherapies or in combination with ATV remained mostly clear of parasites throughout the study and only a few sporadic events of recrudescence were noted. While mice treated with ELQ-331 alone showed no parasitemia during the 45-day monitoring period, in the atovaquone- and ELQ-468-treated groups, two out of ten and two out of five mice, respectively, started showing parasitemia from *ca.* DPI 20 and had to be euthanized by DPI 26 (Fig 7A and 7C). In the case of the combination therapies, two out of ten mice in the ELQ-331 + ATV-treated group developed parasitemia from DPI 24 and had to be euthanized by DPI 26 and one out of ten mice in the ELQ- 468 + ATV-treated group showed parasitemia from DPI 23 and had to be euthanized by DPI 24 (Fig 7B and 7C). As both atovaquone and ELQs are known inhibitors of the parasite cytochrome *bc1* complex, genomic DNA was extracted from recrudescent parasites and the *BdCytb* gene was sequenced. No evidence of mutations was found suggesting that recrudescence is not associated with the development of drug resistance.

**Fig 7.**
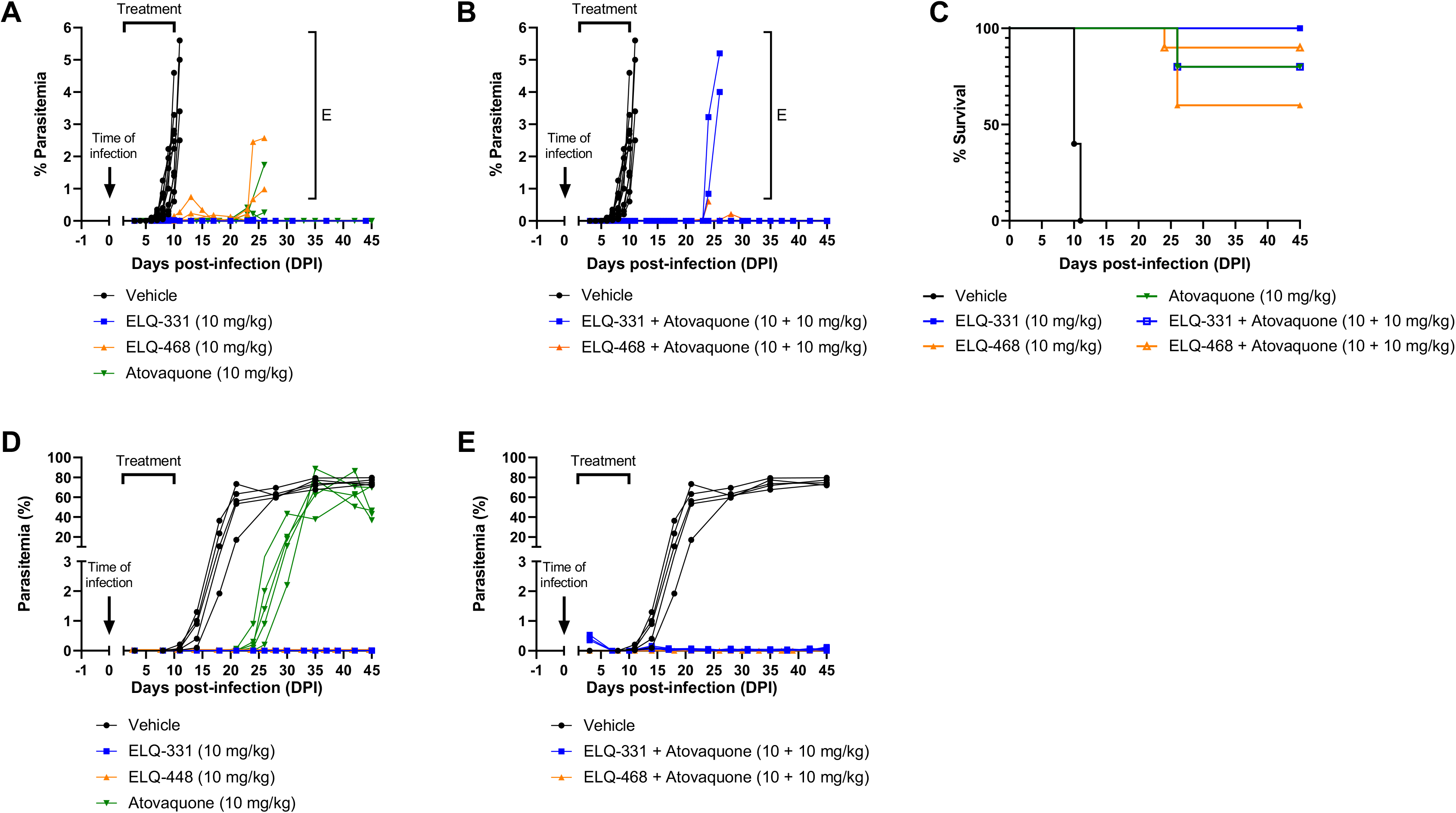
*In vivo* efficacy of ELQ-331 and ELQ-468 against *B. duncani*- and *B. microti*-infected mice. **A-B)** Parasitemia profiles of female C3H/HeJ mice infected IV with 104 *B. duncani*-iRBCs and treated with (**A**) monotherapies of ELQ-331, ELQ-468 or atovaquone at 10 mg/kg or (**B**) combination therapies of ELQ-331 + atovaquone or ELQ-468 + atovaquone at 10 + 10 mg/kg. Treatment was administered daily for 10 days (DPI 1-10) by oral gavage. Control animals received the vehicle alone (PEG-400). E: euthanized. **C)** Kaplan-Meier plot showing percent survival of *B. duncani*-infected mice following treatment with mono- or combination therapies indicated above. **D-E)** Parasitemia profiles of female C.B-17.SCID mice infected with 104 (IV) or 106 (IP) *B. microti*-iRBCs and treated with (**D**) monotherapies of ELQ-331, ELQ-468 or atovaquone at 10 mg/kg or (**E**) combination therapies of ELQ-331 + atovaquone or ELQ-468 + atovaquone at 10 + 10 mg/kg. Treatment was administered daily for 10 days (DPI 1-10) by oral gavage. Control animals received the vehicle alone (PEG-400). E: euthanized.

### *In vivo* efficacy of ELQ-331 and ELQ-468 in *B. microti*-infected mice

The promise delivered by ELQ-331 and ELQ-468 therapies against *B. duncani* infection in mice steered us to evaluate these compounds in a model of *B. microti* infection in SCID mice [23]. Female C.B-17.SCID mice (n = 5 mice/group) were infected with 10^4^ (IV) or 10^6^ (IP) *B. microti*- iRBCs and treated with either monotherapies of ELQ-331, ELQ-468 or atovaquone at 10 mg/kg, or combination therapies of ELQ-331 + atovaquone or ELQ-468 + atovaquone at 10 + 10 mg/kg for 10 days (DPI 1-10). While the vehicle-administered control mice developed high levels of parasitemia by DPI 20 followed by a plateau at 60-80% parasitemia until the end of the study (DPI 45), mice that received ELQ-331 or ELQ-468, alone or in combination with atovaquone, showed no signs of infection during the entire duration of study (Fig 7D and 7E). Mice treated with atovaquone alone showed recrudescence by DPI 24 (Fig 7D). No mutations were found in the *BmCytb* gene of recrudescent parasites.

## Discussion

Here we report the establishment of the in culture-in mouse (ICIM) model of *B. duncani* infection that allows analysis of the intraerythrocytic development of the parasite in human red blood cells *in vitro* and in a mouse model of lethal infection. We further demonstrate that the current clinically approved drug combinations for human babesiosis, azithromycin + atovaquone and quinine + clindamycin are not effective in mice infected with *Babesia* parasites despite showing synergistic interactions *in vitro*. Additionally, we identify two novel compounds, the endochin-like quinolone prodrugs ELQ-331 and ELQ-468, that effectively inhibit *B. duncani* growth *in vitro* at low nanomolar levels and serve as highly effective treatment options against *in vivo* proliferation of both *B. duncani* and *B. microti* parasites.

Over time, our understanding of the basis of life has mostly relied on the use of model organisms and strongly continues to do so [24–27]. In the case of apicomplexan parasites such as *Plasmodium spp*. or *Babesia spp*., the evaluation of novel therapeutics and better understanding of pathogen biology is often challenging because of the lack of strong *in vitro* and *in vivo* models of the same species. For example, research on *P. falciparum*, one of the main causative agents of human malaria, is significantly hindered by the absence of cost-effective and reliable *in vivo* model in small animals. Similarly, research on *B. microti*, the parasite responsible for most cases of human babesiosis, is challenging due to the absence of a continuous *in vitro* culture system. Alternatives such as the use of humanized mice for *P. falciparum* [5, 28, 29] or short-term *ex vivo* assays for *B. microti* [23, 30] have been developed, but they are not ideal for thorough investigation of drug mechanism, host-pathogen interactions or high-throughput screening of chemical libraries. In this regard, *B. duncani*, another agent responsible for human babesiosis, appears to be an ideal candidate for easy *in vitro*-*in vivo* translation. The availability of a continuous *in vitro* culture system in human red blood cells [16] and the ability to propagate this parasite in mice [17, 20, 31] and hamsters [15, 19] offer a complete system for a full characterization of novel therapies and a better understanding of the parasite biology.

Following the development of the *in vitro* culture system of *B. duncani* in human RBCs [16], a key step in the optimization of the ICIM model was to determine whether *B. duncani* infection could be propagated from *in vitro* culture in human erythrocytes to mice. We found that IV injection with either *B. duncani*-infected hRBCs or cell-free merozoites isolated from *in vitro B. duncani* culture led to successful establishment of infection in C3H/HeJ mice. In comparison to mouse-to-mouse transmission of the parasite at an equivalent dose, the delay that is observed in the onset of infection may be attributed to the elimination of much of the parasitized human RBCs by the mouse immune system. Despite the higher infection dose needed to establish a comparable infection profile, the possibility of establishing *B. duncani* infection in mice from an *in vitro* culture offers an interesting alternative to the use of frozen stocks of infected mouse RBCs or the maintenance of live stocks, thus reducing the time and number of animals required to obtain a viable stock.

Another requirement for the establishment of the ICIM model of *B. duncani* was the development of a solid and reliable *in vivo* system of *B. duncani* infection in mice. We investigated the course and severity of *B. duncani* infection in C3H/HeJ mice following the inoculation of different doses of infected RBCs. The observed parasitemia profiles followed that of a typical dose-response study, where the onset of infection, parasitemia levels and severity of the disease increased with the injected dose. The onset of parasitemia was rapid for infection doses of 10^7^ and 10^6^ iRBCs, with levels reaching as high as 35% by DPI 5 and a rapid development of pathology requiring animal euthanasia by DPI 7. Administration of lower doses of iRBCs (10^5^ to 10^2^) resulted in a slow onset of infection with parasitemia levels remaining low (∼ 1-9%). Even with a low parasitemia, all animals ultimately became sick and required euthanasia. In accordance with previous observations [31], we also found that male mice typically develop lower parasitemia than female mice, despite the same fatal endpoint. With regards to the severity of the infection in mice, previous studies suggest that pro-inflammatory cytokines, such as tumor necrosis factor (TNF)-α and interferon (IFN)-γ, play an important role in *B. duncani*-mediated pathogenesis [32, 33]. In some cases, the over-expression of these cytokines can be detrimental to the host and enhance the infection instead of controlling it. Furthermore, IFN-γ and TNF-α appear to interact synergistically during the response to *B. duncani* infection and are likely associated with the pulmonary complications frequently observed in *B. duncani* pathogenesis [32, 33]. Based on these findings, different infection doses could result in different cytokine responses and impact the rapidity and severity of the disease.

To further evaluate the robustness of the *in vivo* model of *B. duncani* infection, we investigated the importance of the route of infection (IP *vs* IV), as well as the genetic background of the host (C3H/HeJ, Balb/cJ and C57BL/6J). As expected, C3H/HeJ mice were readily susceptible to *B. duncani* infection (0% survival) with a rapid onset of parasitemia reaching up to ∼ 10%, whereas C57BL/6J mice appeared resistant to the infection (100% survival) and remained clear of parasitemia throughout the duration of the study (24 days). Although previous studies reported successful *B. duncani* infection in C57BL/6J, the administered dose used in those experiments was much higher than the one used in the present study [21]. Interestingly, Balb/cJ mice showed an intermediate profile between the susceptible C3H/HeJ mice and the resistant C57BL/6J mice, resulting in 40-60% survival rates following *B. duncani* infection. Although in some cases IP infection resulted in a lower peak parasitemia than IV infection, no significant difference was observed in parasitemia profiles or survival following IV or IP administration of the parasites.

To determine whether the apparent resistance of C57BL/6J mice comes from the rapid clearance of the parasite by the immune system or from the inability of the parasite to infect RBCs in this genetic background, we extended the evaluation of *B. duncani* infection to severely immunocompromised mice from the same backgrounds as previously investigated (C3H/HeJ-SCID, Balb/cJ-SCID and C57BL/6J-SCID). All SCID mice became infected, showed significantly higher parasitemia levels than their WT counterparts (66% *vs.* 10% for C3H/HeJ, 60% *vs.* 8% for Balb/cJ and 40% *vs.* 0% for C57BL/6J) and were euthanized by DPI 11-22. It is worth noting that one mouse from the Balb/cJ-SCID cohort cleared the infection following a peak parasitemia around 6%, suggesting a possible leakiness in the *scid* phenotype (Fig 4B). Taken together, these studies indicate that susceptibility to *B. duncani* infection is strongly linked to the host immune response.

To understand whether the stark difference in susceptibility to *B. duncani* infection between C3H/HeJ and C57BL/6J could be attributed to the *tlr4* deficiency reported in C3H/HeJ mice [22], C3H/HeOuJ (WT *Tlr4*) and C57BL/6J(Cg) *Tlr4* KO mice were infected with the same dose of 10^4^ *B. duncani*-iRBCs. The infection profiles in C3H/HeOuJ and C57BL/6J(Cg) *Tlr4* KO mice followed the same trend as their respective parental strain, *i.e.* C3H/HeOuJ mice rapidly developed significant parasitemia and had to be euthanized by DPI 9, whereas C57BL/6J(Cg) *Tlr4* KO mice remained clear of parasitemia throughout the duration of the study. These results indicate that the *tlr4* gene has no apparent effect on the susceptibility to *B. duncani* infection. Further studies are needed to elucidate the possible reasons behind the resistance of certain mouse backgrounds to *B. duncani* infection.

To further validate the *B. duncani* ICIM model, we used it to evaluate the *in vitro* and *in vivo* potency of the drug combinations currently used in the treatment of human babesiosis, as well as two novel endochin-like quinolones, ELQ-331 and ELQ-468, potent antibabesial drugs with low nanomolar efficacy against *B. duncani* [17]. The current arsenal for the treatment of human babesiosis mainly consists of two drug combinations: azithromycin + atovaquone and quinine + clindamycin [34]. These combinations were implemented solely based on their known activity against other apicomplexan parasites, mainly *Plasmodium spp.*, and are frequently associated with severe side effects and/or the development of drug resistance [34]. Recent reports have shown that, except for atovaquone, all three other clinically approved drugs have a weak activity against *B. duncani in vitro* [16]. *In vivo* evaluation of the four drugs yielded similar results, showing high efficacy of atovaquone against *B. duncani* [17] and *B. microti* [17, 23], whereas the other drugs showed no efficacy at doses as high as 50 mg/kg for azithromycin and clindamycin and 100 mg/kg for quinine [35]. Although synergy was observed between the clinically approved drug combinations in *in vitro* assays, no effect was observed against *B. duncani in vivo* following the administration of quinine + clindamycin at 10 + 10 mg/kg and only limited efficacy was noted following treatment with azithromycin + atovaquone at 10 + 10 mg/kg. Despite their similarities with *Plasmodium* parasites, the lack of a food vacuole and/or their inability to degrade hemoglobin might explain why *Babesia spp.* respond poorly to the current therapeutics. Overall, the lack of efficacy of the clinically approved drugs for the treatment of human babesiosis highlights the need for new derivatives specifically designed to tackle *Babesia spp*. infection.

Recent studies identified the prodrugs ELQ-331 and ELQ-468 as potent antibabesial drugs, with IC50 values of 141 nM and 15 nM, respectively [17]. *In* vitro drug efficacy studies using ELQ-300 and ELQ-446, the parent compounds of ELQ-331 and ELQ-468, respectively, showed similar high potency against *B. duncani*, with IC50 values of 36 nM and 34 nM, respectively. Consistent with previous reports investigating other ELQs [17, 35], drug-drug interaction studies showed that ELQ-331 and ELQ-468 are synergistic with atovaquone. *In vivo* evaluation of these derivatives further revealed the high potency of ELQ-331 and ELQ-468 against *B. duncani* as well as *B. microti* infections, both as monotherapies and in combination with atovaquone, highlighting the two ELQ derivatives as promising candidates for the treatment of human babesiosis and *B. duncani* as a suitable model for the identification and evaluation of novel antibabesial therapies.

The continuous *in vitro* culture system of *B. duncani* in human erythrocytes offers a suitable platform to conduct a variety of assays to (a) identify novel and potent antiparasitic drugs (high-throughput screening of chemical libraries, drug efficacy studies, drug-drug interaction studies); (b) understand the mode of action and mechanism of resistance of antiparasitic drugs (selection of drug resistant parasites, propagation of recrudescent parasites from *in vivo* experiments); and (c) unravel the metabolic requirements of parasites for their survival within host red blood cells. Similarly, the *in vivo* model of *B. duncani* infection in mice allows the evaluation of drug efficacy and relapse, the study of virulence and pathology, the investigation of tissue distribution and persistence/dormancy mechanisms, the understanding of host-pathogen interactions and the identification of promising vaccine candidates as well as the evaluation of their protective properties. Furthermore, the ability to propagate *in vitro*-cultured parasites in mice paves the way for a wide variety of applications and the possible development of new animal models using culture-engineered parasites (fluorescent/luminescent parasites, targeted gene deletion to identify genes associated with virulence and pathogenesis, etc.). Finally, the use of *B. duncani* as a model organism could help advance our understanding of the biology of other apicomplexan parasites such as *Plasmodium* and *Toxoplasma spp. B. duncani* could be used as a surrogate to express genes from these parasites, characterize potential vaccine candidates, and unravel the mode of action of antiparasitic drugs.

## Materials and methods

### Chemicals

Atovaquone (A2545, TCI), azithromycin (PHR1088, Sigma), clindamycin (C5269, Sigma), and quinine (22620, Sigma) were purchased from commercial sources. WR99210 was obtained from Jacobus Pharmaceuticals, NJ. *Meta*-chloroperoxybenzoic acid (MCPBA) was purchased from Sigma-Aldrich Chemical Company in St. Louis, MO (USA) and was purified before use according to the method of Armarego and Lin Chai [36]. Analytical TLC utilized Merck 60F-254 250 micron pre-coated silica gel plates and spots were visualized under 254 nm UV light. Flash chromatography over silica gel column was performed using an Isolera One flash chromatography system from Biotage, Uppsala, Sweden. ^1^H-NMR spectra were obtained using a Bruker AMX-400 NMR spectrometer operating at 400.14 MHz. The NMR raw data were analyzed using the iNMR Spectrum Analyst software. ^1^H chemical shifts are reported in parts per million (ppm) relative to internal (TMS) or residual solvent peak. Coupling constant values (J) are reported in hertz (Hz). Decoupled ^19^F operating at 376 MHz was also obtained for compounds containing fluorine (data not shown). High resolution mass spectrometry (LC/MS, HRMS) was performed using a high-resolution (30,000) Thermo LTQ-Orbitrap Discovery hybrid mass spectrometry instrument (San Jose, CA) equipped with an electrospray ionization source operating in the positive or negative mode. The Orbitrap was externally calibrated prior to data acquisition allowing accurate mass measurements for [M+H]^+^ ions to be obtained to within 4 ppm. ELQ-300, ELQ-331, and ELQ-422 were prepared as described previously [37–40]. The alkoxyethyl carbonate ester N-oxide ELQ-468 was synthesized by oxidation of ELQ-422 using *meta*-chloroperoxybenzoic acid (MCPBA) as shown in the Scheme 1.

**Scheme 1.**
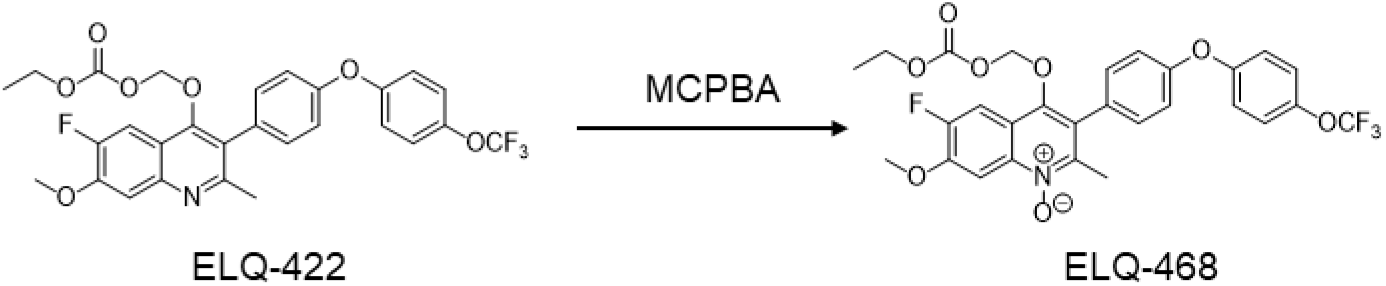
Chemical Synthesis of ELQ-468.

### 4-(((ethoxycarbonyl)oxy)methoxy)-6-fluoro-7-methoxy-2-methyl-3-(4-(4-(trifluoromethoxy)-phenoxy)phenyl)quinoline 1-oxide (ELQ-468)

To a stirred solution of ELQ-422 (1.12 g, 2.0 mmol) in chloroform (50 mL) was added MCPBA (692 mg, 4.0 mmol) and heated at 80°C for 24 hours. After cooling to room temperature, the yellow solution was roto-evaporated under vacuum to dryness and the product was purified by flash chromatography twice using a gradient of ethyl acetate/methylene chloride as eluent to afford ELQ-468 (847 mg, 73.6 % yield) of a white oil, which solidified upon standing at room temperature as a white solid. ^1^H-NMR (600 MHz; CDCl_3_): δ 8.29 (d, *J* = 7.9 Hz, 1H), 7.73 (d, *J* = 11.0 Hz, 1H), 7.38-7.36 (m, 2H), 7.28 (d, *J* = 8.1 Hz, 3H including the residual proton from CDCl_3_), 7.16-7.13 (m, 4H), 5.26 (s, 2H), 4.13-4.10 (m, 5H), 2.57 (s, 3H). High-resolution mass spectrum calculated for [M+H]_+_ = 578.1433, observed for [M+H]_+_ = 578.1440.

### In vitro B. duncani culture

*In vitro* propagation of *B. duncani* in human red blood cells (hRBCs) was carried out as previously reported by Abraham *et al.* [16] and Chiu *et al.* [17]. Briefly, cryo-preserved *Babesia duncani* WA-1 strain [14] was thawed as per standard method [17]. *B. duncani* parasites were maintained in A^+^ hRBCs (American Red Cross) at 5% hematocrit in HL-1 base medium (Lonza 344017) supplemented with 20% heat-inactivated fetal bovine serum (FBS; Sigma F4135), 2% 50X HT media supplement Hybri-max (Sigma H0137), 1% of 200 mM L-glutamine (GIBCO 25030-081), 1% 100X penicillin-streptomycin (GIBCO 15240-062) and 1% 10 mg/mL gentamicin (GIBCO 15710-072). Additionally, *in vitro* parasite propagation was also conducted in Claycomb medium (Sigma 51800C) supplemented as described above. Parasite cultures were incubated at 37°C under 2% O2, 5% CO2 and 93% N2 gas mixture in a humidified chamber. Evolution of parasitemia was monitored either by light microscopy examination of Giemsa-stained blood smears or by fluorescence detection of SYBR Green I, following a previously reported method [16]. Briefly, a 1:1 mixture of parasite culture and lysis buffer (0.008% saponin, 0.08% Triton-X-100, 20 mM Tris-HCl (pH 7.5) and 5 mM EDTA) containing SYBR Green I (0.01%) was incubated at 37°C for 1 h in the dark. Fluorescence was then measured for λex = 497 nm and λem = 520 nm using a BioTek Synergy™ Mx Microplate Reader. Background fluorescence from uninfected RBCs was subtracted from each measurement.

### Purification of *B. duncani* merozoites

*B. duncani* WA-1 parasites were cultured to 18-20% parasitemia and blood smears were used to confirm the presence of free merozoites. High parasitemia culture containing a large number of free merozoites was centrifuged at 1800 rpm for 5 min at 37°C. The supernatant containing free merozoites was collected and centrifuged at 4000 rpm for 10 min at 37°C. The pellet containing free merozoites was mixed with warm Claycomb medium in 1:5 (w/v) ratio. Purity of the merozoite suspension was assessed by light microscopy examination of Giemsa-stained blood smears.

### *In vitro* drug efficacy assays

The effect of anti-parasitic drugs on the parasite’s intraerythrocytic developmental cycle (IDC) was evaluated by treating parasites with a gradient of drug concentrations and determining the median inhibitory concentration (IC50) following a previously established protocol [17]. Briefly, *B. duncani in vitro* culture (0.1% parasitemia) in 5% hematocrit was treated for 60 h with concentrations starting at 10 µM and 2-fold serially diluted. Treatment with 2 µM WR99210 was used as positive control, representing 0% growth and untreated iRBCs were used as negative control, representing 100% growth. Following treatment, parasitemia was quantified using SYBR Green I, as described above. Fluorescence readings were normalized to the positive and negative control values. IC50 values were determined from a sigmoidal dose-response curve using GraphPad Prism version 9.2.1. Data are presented as mean ± SD of two independent experiments performed in biological triplicates.

### *In vitro* drug-drug interaction assays

A modified fixed ratio method of *in vitro* drug combination assay [40] was used to understand the type of interaction between two drugs of interest against *B. duncani* infection. Briefly, drug combinations were used in four fixed ratios (4:1, 3:2, 2:3 and 1:4) with an initial concentration of each drug fixed at approximately 8X IC50. These drug combinations were serially diluted (2-fold) six times so that the IC50 concentration of an individual drug falls within the 3^rd^ and 4^th^ dilutions. Parasites (0.1% parasitemia, 5% hematocrit, 200 μL) were incubated with each drug combination for 60 h at 37°C. Parasitemia was determined using SYBR Green I as described above. The mean fractional inhibitory concentration (FIC50) for each drug was determined using the following equation: FIC50 = (IC50 of the drug in combination) / (IC50 of drug alone). The interaction between two drugs A and B was determined by the sum of FIC50 (ƩFIC50) using the following formula: ƩFIC50 = [(IC50 of drug A in combination) / (IC50 of drug A alone) + (IC50 of drug B in combination) / (IC50 of drug B alone)]. Isobolograms were plotted as FIC(drug A) = f(FIC(drug B)). Drug combinations with ƩFIC values ≤ 0.8 represent drugs interacting in a synergistic manner, while drug combinations with ƩFIC ≥ 1.4 represent antagonistic interactions [41].

### *In vitro* toxicity assay in mammalian cells

Drug activity in mammalian cells was evaluated in HeLa, HepG2 and HCT116 cells as described by Chiu *et al.* [17].

### Ethics Statement

All animal experiments were approved by the Institutional Animal Care and Use Committees (IACUC) at Yale University (Protocol #2020-07689). Animals were monitored for one week after arrival before the start of an experiment. Animals that showed signs of distress or appeared moribund were humanly euthanized using approved protocols.

### In vivo B. duncani infection

Five-to six-week-old female and male mice of the following strains were purchased from Jackson Laboratories: C3H/HeJ, Balb/cJ, C57BL/6J, C3H/HeOuJ, C57BL/6J(Cg) *Tlr4* KO, C3H/HeJ-SCID, Balb/cJ-SCID and C57BL/6J-SCID. *B. duncani* (WA-1) parasites were thawed from previously cryo-preserved infected mouse blood and were subsequently propagated in C3H/HeJ mice. Mice were inoculated with either blood from an infected stock mouse at indicated doses, *in vitro*-cultured infected hRBCs (8.5 x 10^5^) or purified merozoites (2 x 10^7^) *via* intravenous (IV) or intraperitoneal (IP) injection and monitored for establishment of parasitemia. Animals were bled at specified time points either by retro-orbital or tail vein bleeding and parasitemia was determined by light microscopy examination of Giemsa-stained thin blood smears.

### In vivo B. microti infection

Five- to six-week-old female C.B-17.SCID (C.B-17/IcrHsd-Prkdc^scid^) mice were obtained from Envigo. *B. microti* (LabS1) parasites were thawed from previously cryo-preserved infected mouse blood and were subsequently propagated in C.B-17.SCID.

### *In vivo* drug efficacy in *B. duncani-* and *B. microti-* infected mice

Six- to seven-week-old female C3H/HeJ mice (n = 5-10 mice/group) were infected IV with 10^3^ *B. duncani*-infected mouse RBCs (mRBCs). Mice were treated by oral gavage (OG, 100 μL) for 10 days (DPI 1-10) with the vehicle (PEG-400), ELQ-331, ELQ-468 or atovaquone alone at 10 mg/kg, or with the following combinations: azithromycin + atovaquone, quinine + clindamycin, ELQ-331 + atovaquone or ELQ-468 + atovaquone at 10 + 10 mg/kg.

Six- to seven- week-old female C.B-17.SCID mice (n = 5 mice/group) were infected with either 10^4^ (IV) or 10^6^ (IP) *B. microti*-infected mRBCs. Mice were treated by oral gavage (OG, 100 μL) for 10 days (DPI 1-10) with the vehicle (PEG-400), ELQ-331, ELQ-468 or atovaquone alone at 10 mg/kg, or with the following combinations: ELQ-331 + atovaquone or ELQ-468 + atovaquone at 10 + 10 mg/kg. Parasitemia was determined by light microscopy examination of Giemsa-stained thin blood smears.

## Acknowledgements

The authors would like to thank Heather Wallace from Yale Animal Resources Center, for performing oral gavage in *in vivo* studies.

